# Development of a 3D *in vitro* wound healing model to assess the effect of ADSC-EVs on vascularisation

**DOI:** 10.1101/2025.07.16.665214

**Authors:** Emma K. C. Symonds, Alfonso J Schmidt, Alexander W Brown, Margaret J Currie, Patries M Herst, Kathryn E Hally, Kirsty M Danielson

## Abstract

Angiogenesis is critical for effective wound healing and relies on the successful coordination of various cell types including endothelial cells, macrophages, and fibroblasts. Adipose-derived stem cell extracellular vesicles (ADSC-EVs) have demonstrated pro-angiogenic properties and have been posited as a novel therapeutic to aid wound healing; however, their functional impact within human-derived multicellular models remains largely uncharacterised. This study explores the development and application of a 3D multicellular *in vitro* model to assess the effects of ADSC-EVs on vascularisation in the context of wound healing.

3D multicellular *in vitro* models were developed by co-culturing human umbilical vein endothelial cells (HUVECs), monocyte-derived macrophages (MDMs), and fibroblasts within Matrigel to recapitulate the *in vivo* wound healing microenvironment. A five-colour confocal microscopy panel was developed to visualise each cell type and EVs within the models. The optimised models were then treated ADSC-EVs or control to determine their impact on angiogenesis and cell co-localisation.

We determined that vessel formation was significantly enhanced when HUVECs were co-cultured in multicellular models compared to monocultures, with the greatest effect observed in the full three-cell-type model. This effect was even more pronounced with the addition of ADSC-EVs. ADSC-EV treatment also enhanced macrophage co-localisation within endothelial structures.

This study developed a multicellular model that can be used for future work assessing wound healing *in vitro* and will be additive to currently used single-cell and *in vivo* models. We have applied these models to demonstrate that ADSC-EVs significantly enhance tube formation in HUVECs, and the development of tissue-like structures in multicell systems, highlighting their potential as a promising therapeutic approach for improving wound healing.

## Introduction

Wound healing is a complex process that involves tissue repair and regeneration following injury. One of the critical factors in successful wound healing is vascularisation, the formation of new blood vessels at the wound site ^1,2^. Vascularisation is essential for providing oxygen, nutrients, and immune cells to the damaged tissue ^1^. It also plays an important role in the formation of granulation tissue, collagen deposition, and extracellular matrix (ECM) synthesis, all of which are necessary for the structural integrity and restoration of normal tissue architecture. Without adequate vascular supply, healing can be delayed or impaired, leading to chronic wounds or infection ^1-4^.

One promising therapeutic approach in promoting vascularisation and improving wound healing involves the use of mesenchymal stem cell (MSC)-derived extracellular vesicles (EVs). MSC-EVs, particularly those derived from adipose-derived stem cells (ADSCs), have gained significant attention for their ability to modulate key processes in tissue repair, including angiogenesis and inflammatory regulation ^5^. Studies have demonstrated that MSC-EVs and ADSC-EVs can stimulate the expression of angiogenesis-related genes such as ERK, STAT3, and AKT in human dermal fibroblasts and endothelial cells, including human umbilical vein endothelial cells (HUVECs) ^5,6^. Moreover, ADSC-EVs have been shown to regulate the inflammatory environment by modulating cytokines such as interleukin (IL)-1β, interferon (IFN)-*γ*, and tumour necrosis factor (TNF)-α, thus influencing the immune response in wound healing ^7^. We have previously demonstrated that ADSC-EVs reduce the overall inflammatory profile of macrophages, promoting differentiation towards a more M2-like phenotype (alternatively activated macrophages) ^8^. This anti-inflammatory macrophage phenotype is desired in wound healing to minimise inflammation and promote tissue survival.

Despite these promising findings, current research primarily focuses on assessing the bioactivity of EVs using single-cell-type *in vitro* models. These models, often centred on HUVECs, fail to capture the complexity of the wound healing process in which multiple cell types including fibroblasts, macrophages, and endothelial cells, interact within a dynamic microenvironment. Fibroblasts are integral to ECM development and the structural organisation of the wound site. Upon activation by growth factors such as TGFβ-1, fibroblasts differentiate into myofibroblasts, which play a pivotal role in wound contraction and tissue remodelling ^9-11^. Following tissue injury, macrophages are also recruited to the site and contribute to angiogenesis by releasing growth factors that induce capillary sprouting and vessel formation around the ECM scaffold established by fibroblasts ^12,13^.

Given the importance of these cellular interactions in vascularisation and wound healing, there is a clear need for more sophisticated, multi-cell-type *in vitro* models that can better recapitulate the complexity of the wound healing microenvironment. Such models will not only provide a more accurate representation of the cellular dynamics of wound healing but also enhance our understanding of how ADSC-EVs can be harnessed as acellular therapeutics to improve wound healing outcomes. This study develops and applies a multi-cell-type *in vitro* model to study the effects of ADSC-EVs on vascularisation during wound healing, with the aim of advancing therapeutic strategies for effective tissue regeneration.

## Methods

### Participant Recruitment, ADSC isolation, and extracellular vesicle isolation

ADSCs were isolated from the lipoaspirate of three participants recruited as part of a larger ongoing study on autologous fat graft retention. This study has ethical approval from the Health and Disabilities Ethics Committee, New Zealand (19/CEN/23) and all participants provided written, informed consent. Adipose tissue samples were collected at the time of surgery and transferred to a sterile facility. Cells were isolated using 0.1% Type IA Collagenase (Sigma-Aldrich, MO, USA) and cultured at 37^°^C with 5% CO_2_. We have previously confirmed isolated cells as ADSCs according to International Society for Cell Therapy guidelines ^8^. Conditioned culture media (DMEM + 10% EV-depleted FBS + 1% penicillin-streptomycin) was collected at passages 2-5, centrifuged at 2000 x g for 10 min, and slow frozen using isopropanol to -80°C for downstream EV isolations. To generate EV-depleted FBS, FBS was diluted 1:4 in culture media, ultracentrifuged at 100 000 x g for 16 hrs, and the supernatant was collected and stored at -80°C until required.

We have submitted all relevant data from our experiments to the EV-TRACK knowledgebase (EV-TRACK ID: EV230010) ^14^. For EV isolations, cultured media supernatants were thawed at 37^°^C, centrifuged at 2000 x g for 5 min, and EVs were isolated from 10 mL of supernatant using a combination of size exclusion chromatography (SEC) and ultrafiltration. In brief, EVs were separated using SEC columns (qEV10/35nm) on an automated fraction collector (Izon, New Zealand), fractions 1-4 were collected and pooled, and the total volume was concentrated using Amicon Ultra-15 Centrifugal Filter Units by centrifuging at 4696 x g for 35 min. Dummy-EVs (D-EVs) were defined as cell culture media that was cultured in the absence of ADSCs and underwent the same isolation process (SEC and ultrafiltration) as media cultured in the presence of ADSCs. This isolate was used as a control for any non-EV particles potentially arising from the culture media, and any remaining FBS-derived EVs. ADSC-EVs were previously characterised using tunable resistive pulse sensing, Western blot, and transmission electron microscopy ^8^. 1×10^5^ particles/mL were added to downstream cell culture assays, based on a previous dose response experiment ^8^, and an equal volume of D-EVs was added to control wells. For all experiments, ADSC-EVs from three patients, or D-EVs, were added to the media (each patient added to a separate well) with each group seeded in duplicate and incubated at 37°C/5%CO_2_.

### HUVEC culture

HUVECS (PCS-100-013, ATCC) were used as a baseline single-cell-type model of angiogenesis for comparison to multicellular models. 300 μL of Matrigel (Corning Life Sciences, NY, USA) was used to coat the wells of 24 well plates. HUVECs (passage 3) were resuspended in media (Vascular Basal Cell Medium + 2% EV-depleted FBS + 1% penicillin streptomycin + Endothelial Cell Growth Kit, ATCC) at 1×10^5^ cells/mL and incubated with 1% Calcein AM for 30 min at room temperature. Cells were pelleted and resuspended to remove excess stain, and 500 μL of the cell suspension was added on top of the pre-set Matrigel. ADSC-EVs were added to the media at a concentration of 1×10^5^ particles/mL, with an equal volume of D-EVs used for control wells. Cultures were incubated at 37°C/5%CO_2_ for 24 hrs before visualisation on a fluorescent microscope (CKX53, Olympus, Tokyo, Japan). Tube formation was assessed using AngioTool software (version 6.0) ^15^.

### Monocyte derived macrophage (MDM) isolation and culture

Self-reporting healthy volunteers were recruited and provided written, informed consent (HDEC 19/CEN/129) for venepuncture. Peripheral blood mononuclear cells (PBMCs) were isolated from the buffy coat of 20 mL of whole blood using RosetteSep human enrichment cocktail (STEMCELL Technologies, BC, Canada) and density gradient centrifugation (Ficoll-Paque PLUS, Bio-Strategy, Australia) as per the manufacturer’s instructions. Isolated cells were cultured in RPMI + 10% FBS + 1% penicillin-streptomycin (P-S; In Vitro Technologies) at 37°C/5%CO_2_ for 24 hrs. On day one, media was changed to M0 (undifferentiated macrophages) stimulating media (RPMI + 10% EV-depleted FBS + 1% penicillin streptomycin + 100 ng/mL Macrophage Colony Stimulating Factor).

### Development of 3D multicellular models incorporating two cell types

A four-colour confocal microscopy panel was designed to assess 3D multicellular models that incorporate two cell types. Initially, two models were developed; Model 1 (HUVECs + MDMs + ADSC-EVs) and Model 2 (HUVECs + fibroblasts + ADSC-EVs). The confocal panel is outlined in Table 1 and model development is outlined in Figure 1.

**Table 1:** Five colour confocal panel for multicellular models.

| Stain | Excitation/Emission | Laser line | Use | Dilution |
| --- | --- | --- | --- | --- |
| NucBlue | 360/460 | 405 | Nuclear | 1 drop/well |
| HLADR-BV480 | 436/478 | 445 | MDMs | 1:200 |
| Calcein AM | 488/520 | 488 | HUVECs | 1:1000 |
| PKH26 | 551/567 | 561 | ADSC-EVs | 1:500 |
| Cell Tracker Deep Red | 630/650 | 640 | Fibroblasts | 1:20000 |

**Figure 1:**
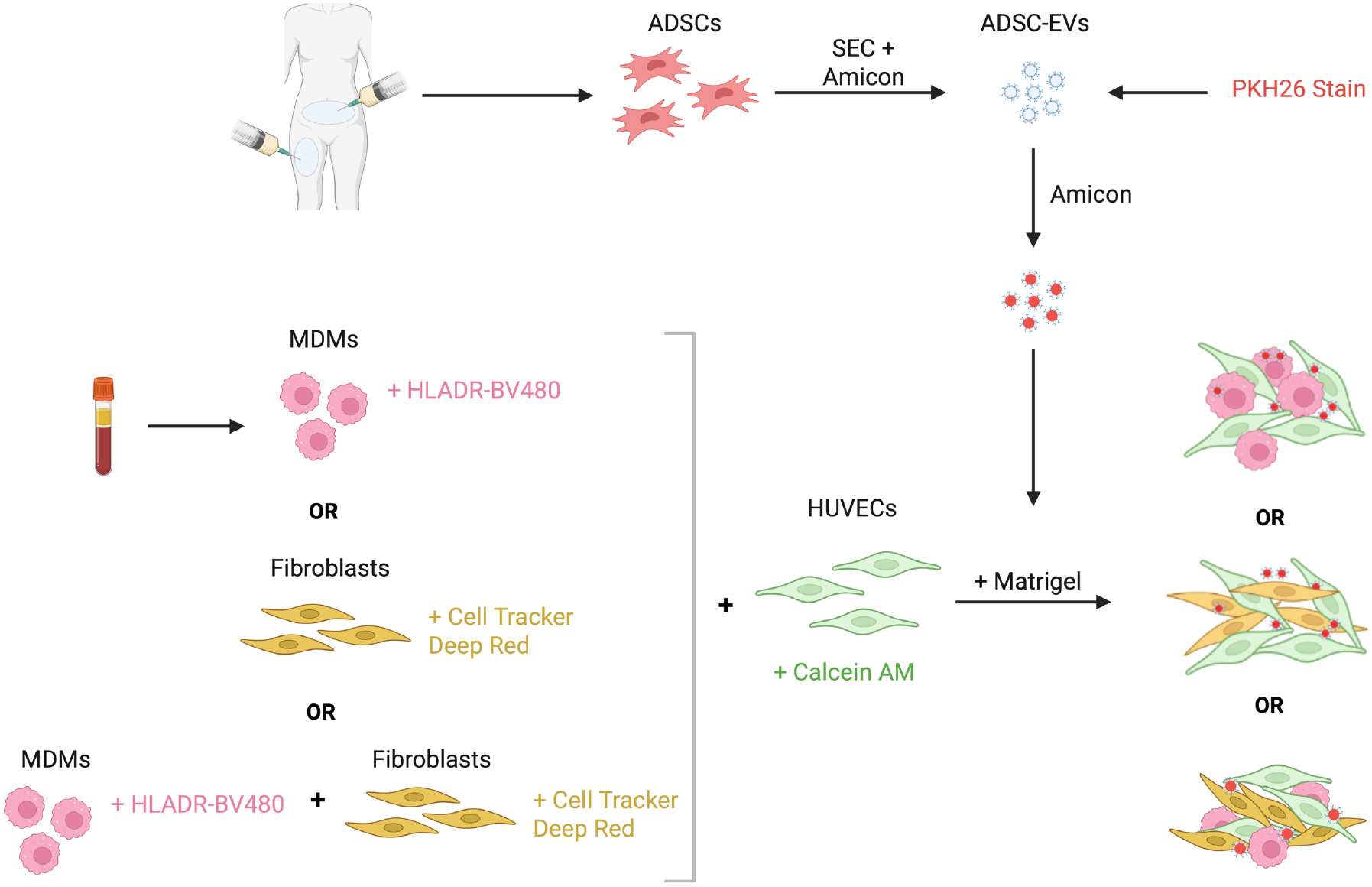
Summary of the development of the multicellular models. Created with BioRender.com.

### MDM and HUVEC model development

Unpolarised M0-like MDMs and HUVECs (passage 3) were cultured together to assess different seeding methodologies within the Matrigel. MDMs from 20 mL of whole blood from a healthy volunteer (n=1) were isolated and cultured as previously described. On day 7, they were lifted from plates using Accutase (Sigma) and incubated in 0.05% Cell Tracker Deep Red (diluted in PBS) at 37°C with 5% CO_2_ for 30 min in the dark. HUVECs were cultured for three days in Vascular Basal Cell Medium + 2% EV-depleted FBS + 1% penicillin streptomycin + Endothelial Cell Growth Kit (ATCC), then incubated in 1% Calcein AM (diluted in media) and cells were incubated for 30 min at room temperature. Three different protocols were then tested to develop the 3D multicellular models. Firstly (Method 1), MDMs and HUVECs (1×10^5^ cells/mL for each cell type) were combined and resuspended in Matrigel before coating wells of a 24-well plate with 150 μL of Matrigel-cell solution. Cultures were then placed in the incubator for 20 min to set before 500 μL of media was added. Secondly (Method 2), MDMs alone (1×10^5^ cells/mL) were mixed in Matrigel, plated, and set for 20 min. HUVECs were resuspended in media at 1×10^5^ cells/mL and 500 μL was added on top of the Matrigel. Lastly (Method 3), wells were coated with Matrigel alone and set. HUVECs and MDMs (1×10^5^ cells/mL for each cell type) were combined and resuspended in media, and 500 μL of the suspension was then added to the top of the Matrigel. Cultures were incubated for 24 hrs at 37°C with 5% CO_2_. One drop of NucBlue was added to each well and incubated at RT in the dark for 30 min. Following analysis, Method 3 was used for all downstream model development and analysis.

### Fibroblast and HUVEC model development

Human breast skin fibroblasts (CCD-1128SK, ATCC; passage 4) were cultured in IMDM + 10% EV-depleted FBS + 1% penicillin streptomycin for three days before being lifted from plates using 0.25% Trypsin (Sigma) and centrifuged at 150 x g for 5 min. Cells were diluted to 1×10^5^ cells/mL and 1 μL of Cell Tracker Deep Red was added for every 1 mL of media. Fibroblasts were incubated at 37°C with 5% CO_2_ for 30 min in the dark. HUVECs were isolated and stained with Calcein AM as described above. Both fibroblasts and HUVECs were centrifuged at 150 x g for 5 min to pellet the cells and remove excess stain. Models were formed and incubated as described above using Method 3.

### MDM, fibroblast, and HUVEC model development

Following the successful development of two-cell-type models, a 3D model was developed using all three cell types. HUVECs, fibroblasts, and MDMs cultured as described above. Wells of a 24 well plate were coated with 150 μL of Matrigel and incubated at 37°C for 20 min to set. HUVECs and fibroblasts were stained with Calcein AM and Cell Tracker Deep Red as described above. MDMs were incubated with HLA-DR-BV480 (1:200 diluted in FACS buffer) for 30 min at 4°C before pelleting and resuspending in culture media. HUVEC, fibroblast, and MDM cell suspensions were combined at a concentration of 1×10^5^ cells/mL of each cell type. 500 μL of the cell suspension was added to each well and incubated for 24 hrs at 37°C with 5% CO_2_. One drop of Nuclear Blue was added to each well and incubated at RT in the dark for 30 min.

### PKH26 optimisation for ADSC-EV visualisation

PKH26 staining of EVs was optimised by comparing stained ADSC-EVs, D-EVs, or detergent treated (Triton X-100; Thermo Fisher Scientific) ADSC-EVs. ADSC-EVs or D-EVs were isolated from cell culture media using SEC columns and Amicon Ultra-15 centrifugal filter units as outlined above. ADSC-EVs were stained using the PKH26 Red Fluorescent Cell Linked Mini Kit (Sigma). Concentrated products of ADSC-EVs or D-EVs were added to 500 μL of Diluent C before being mixed with 2 μL of PKH26 stain that was previously diluted in 500 μL of Diluent C. For the Triton X-100 treated samples, 0.1% Triton X-100 was added to lyse the particles within the concentrated product. All samples were incubated in the dark at RT for 5 min, diluted in PBS to 10 mL, and centrifuged at 4696 x g for 35 min in Amicon Ultra-15 centrifugal filter units to remove excess stain. ADSC-EVs, D-EVs, or Triton X-100 treated ADSC-EVs were then added to the media of HUVEC and MDM cultures, at the same time as the cells themselves, at a concentration of 1×10^5^ particles/mL. Cultures were incubated for 24 hrs at 37°C with 5% CO_2_. One drop of Nuclear Blue was added to each well and incubated at RT in the dark for 30 min.

### Imaging and image analysis

All cultures were visualised and imaged on inverted microscope IX83 equipped with Laser Scanning Confocal Microscope head FV3000 (Olympus, Japan) using software FV31-SW, version 2.4198. The imaging process was done using lasers with 405 nm (50 mW), 445 nm (75 mW) 488 nm (20 mW), 561nm (20 mW) and 640 nm (40 mW) excitation wavelengths. The emission from each fluorophore was collected by highly sensitive detectors configured with emission bandwidth of 430-470, 500-540, 460-500, 570-620, 650-750 respectively. The images were recorded using an UPLXAPO 10X NA 0.4 and UPLXAPO 20X NA 0.75 objectives covering 140 μm Z axis. Images were recorded at 0.7 megapixels with lateral resolution of 1.59 μm and axial resolution of 3 μm. Images were imported to ImageJ (FIJI) ^16^ and processed by applying max Intensity Z projections to obtain an even two dimensional image with the aim visualise the presence or absence of PKH26 staining and further analysis. Additionally, IMARIS software (Oxford Instruments, version 10.2) was used to create 3D visualisations of the models.

### Assessing functional changes in 3D multicellular models with ADSC-EV co-culture

Cell cultures were generated for all three of the developed models (HUVEC/MDM, HUVEC/fibroblast, and HUVEC/MDM/fibroblast) as described above. ADSC-EVs from three consecutive patients, or D-EVs, were isolated and stained as previously described. 1×10^5^ EVs/mL were added to desired cultures at the same time as the cells, with each ADSC-EV patient (n=3) or D-EV sample cultured in duplicate. Models were incubated for 24 hrs. One drop of Nuclear Blue was added to each well 30 min before imaging. Functional changes with the addition of ADSC-EVs were characterised by HUVEC tube formation, as described below.

### In vitro angiogenesis assessed by HUVEC tube formation

Tube formation assays were used to assess the ability of HUVECs to form capillary-like structures in 3D multicellular models. Angiotool software was used to analyse the number of junctions, vessel length, junction density, and vessel percentage area. Tube formation was compared for each of the 3D multicellular models plus ADSC-EVs or D-EVs. This data was compared to HUVEC tube formation in the single-cell-type cultures.

### Assessment of cellular co-localisation

Z projections were further analysed to assess the co-localisation of MDMs or fibroblasts to HUVECs. When fibroblast/HUVEC images were observed, these cells were completely co-located with HUVECs and further analysis was not necessary. To analyse MDM and HUVEC co-localisation, all channels were turned off except for HUVECs (green) and a trace was manually drawn around the HUVEC tubes on Fiji. This channel was then turned off and the channel fluorescing MDMs (magenta) was turned on. The number of MDMs inside the trace was compared to the total number of MDMs visualised in the image, calculated using Automated Cell Counting on Fiji. This was done by converting images to 16-bit grayscale and utilising the “Analyse Particles” feature.

### Statistical Analysis

Fields of view and duplicates were averaged per ADSC-EV sample (n=3), and data are presented as mean ± SD. For the angiogenesis assay, data were graphed to compare tube formation when HUVECS were cultured with either MDMs, fibroblasts, or both, with ADSC-EVs or D-EVs. Co-localisation analysis was normalised to the original HUVEC trace area to account for any changes in tube formation between ADSC-EV and D-EV treated models. Groups were compared using a one-way ANOVA and Tukey’s multiple comparisons tests. All data were analysed using GraphPad Prism (version 9.1.0) and *p*<0.05 was considered statistically significant.

## Results

### Optimisation of cell seeding using HUVEC and MDM model

To optimise the seeding of HUVECs and MDMs, 3D multicellular models were created using three different methods. Firstly, MDMs and HUVECs were both mixed in Matrigel prior to plating, resulting in a lack of tube formation and cell association (Figure 2A). Next, MDMs were mixed in Matrigel and HUVECs were seeded on top. This resulted in successful tube formation, but no interaction between the two cell types as the MDMs spread throughout the Matrigel structure more extensively than the HUVECs (Figure 2B). Lastly, both MDMs and HUVECs were seeded on top of the Matrigel. This method demonstrated tube formation and interaction between the two cell types (Figure 2C). The last method was then used to create a second two-cell-type model (HUVECs/fibroblasts) as well as a three-cell-type model (HUVECs/MDMs/fibroblasts; Figure 2D). To assess the effectiveness and practicality of the developed models, they were subsequently used for downstream functional assays.

**Figure 2:**
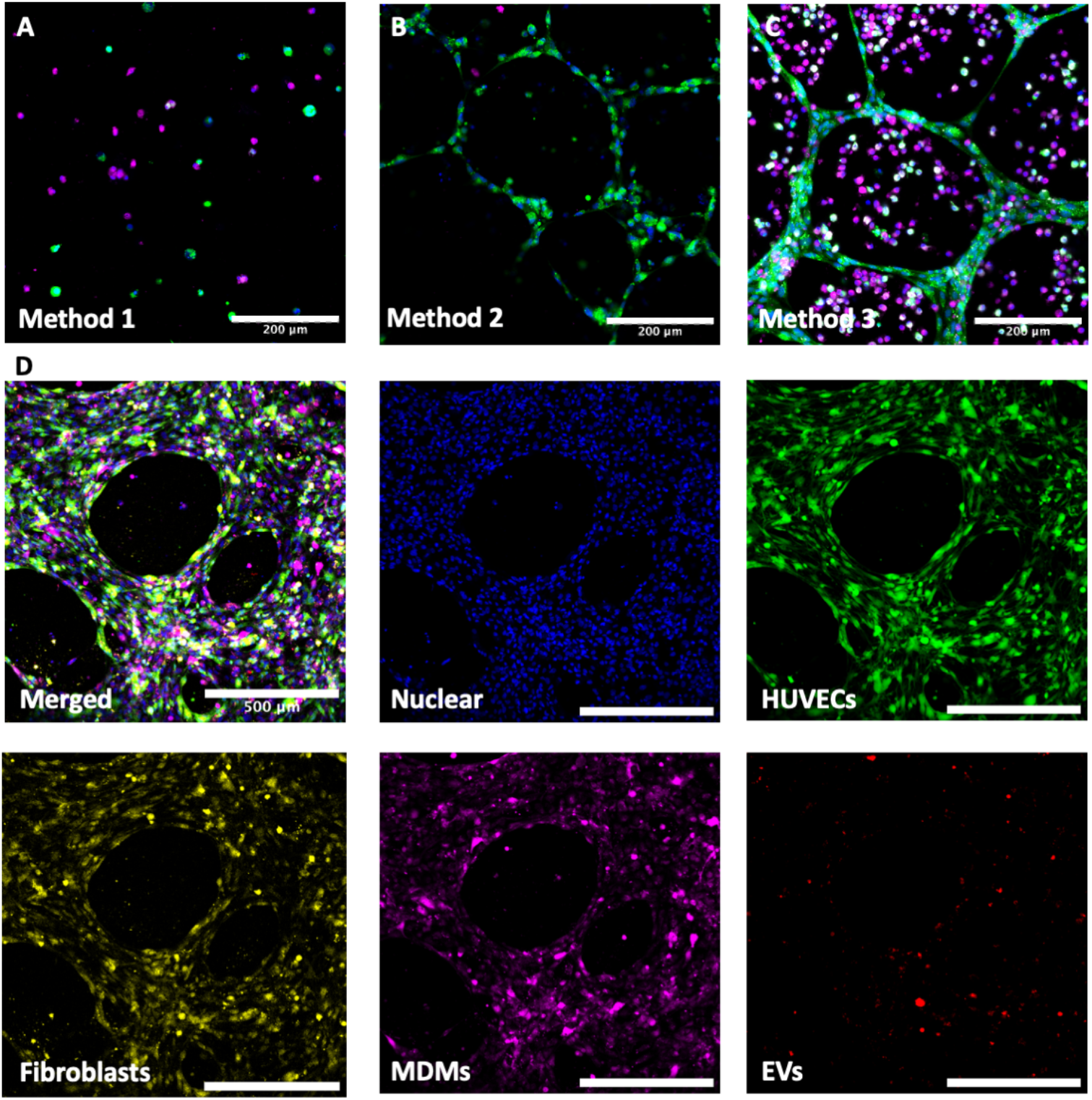
Development of cell culture models. A) To optimise the seeding of MDMs and HUVECs into a Matrigel model, cells were first both mixed with Matrigel, resulting in no tube formation or cell association. B) MDMs were mixed in Matrigel and HUVECs were seeded on top, resulting in tube formation, but no cell association. C) Matrigel was plated first and both cell types were added on top, resulting in tube formation and cell co-localisation. D) Representative images of all individual channels of the complete model. Green = HUVECs, magenta = MDMs, and blue = nuclear stain. White indicates the overlapping of colours. Images were recorded on a Confocal FV3000 Laser Scanning Microscope (Olympus) and Z projections were created using Fiji. Scale bar represents 200 μm (A-C) or 500 μm (D).

### Tube formation is enhanced when HUVECs are co-cultured in multicellular models

HUVEC tube formation was compared between single cell type HUVEC models, HUVEC/MDM models, HUVEC/fibroblast models, and full cell models to determine if the presence of different cell types influenced tube formation. Relative to single cell type cultures (set to 1), co-culture with either MDMs or fibroblasts significantly increased vessel length (macrophages 24.12±5.52, *p*=0.0001, fibroblasts 14.18±2.11, *p*=0.0052 Figure 3C) and vessel percentage area (macrophages 2.38±0.16, *p*=0.0184, fibroblasts 2.79±0.62, *p*=0.0041, Figure 3D). When all three cell types were cultured together, there was a significant increase in vessel length (28.65±2.89, *p*<0.0001, Figure 3C), and vessel percentage area (2.91±0.57, p=0.0027, Figure 3D) compared to HUVECs cultured alone.

**Figure 3:**
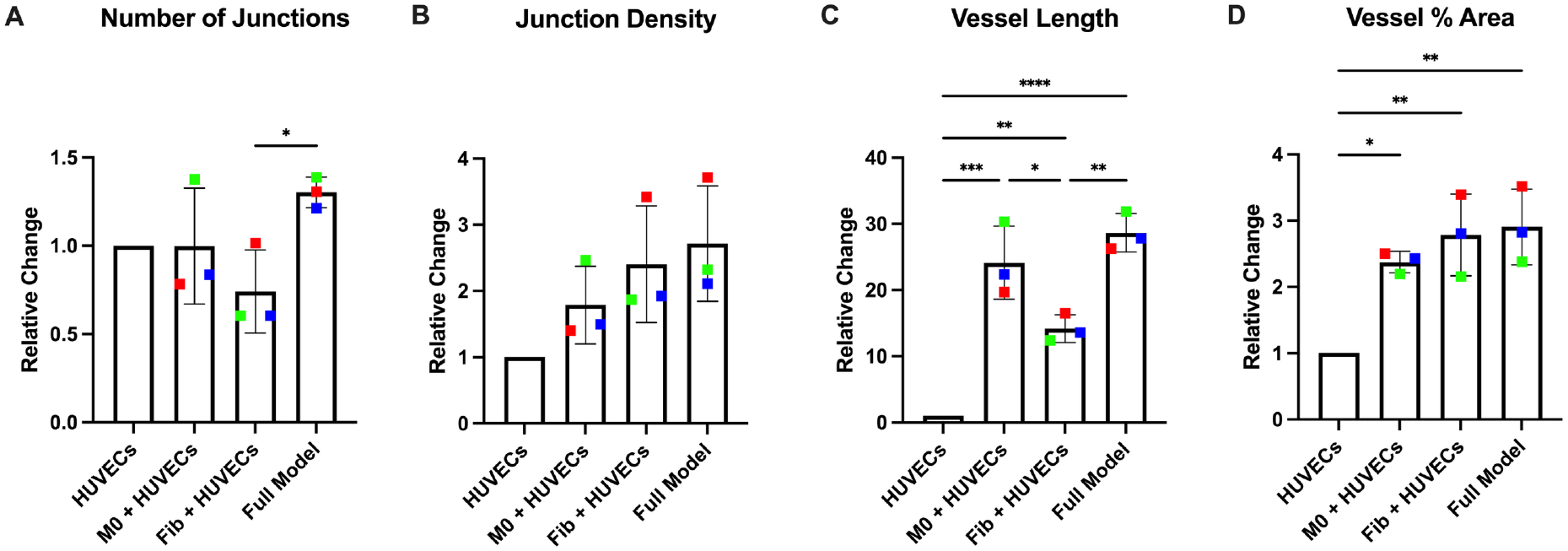
Culturing cells in multicellular models increases tube formation. HUVECs were cultured alone, with MDMs, with fibroblasts, or all three cell types together, to assess tube formation. Groups were compared using a one-way ANOVA and Tukey’s multiple comparisons tests. Coloured dots represent three ADSC patients across 2 experimental runs (averaged). Data were analysed using GraphPad Prism (9.1.0). \**p*<0.05, \*\**p*<0.01, \*\*\*\**p*<0.0001.

### ADSC-EVs are successfully stained and taken up by endothelial cells, MDMs, and fibroblasts in 3D models

To determine if PKH26 successfully and specifically stained ADSC-EVs, the fluorescence of PKH26 was compared in D-EV, ADSC-EV, and Triton X-100 treated ADSC-EV samples that were added to pre-made HUVEC/MDM 3D multicellular models. Very little fluorescence was observed in D-EVs models, and almost no fluorescence was observed in Triton X-100 treated ADSC-EV samples (Figure 4A), indicating that there were very low levels of non-specific staining. In contrast to this, ADSC-EV treated samples demonstrated a high level of fluorescence, illustrated by red staining (Figure 4A). In a further investigation utilising both types of two-cell-type 3D multicellular models (HUVEC/MDMs and HUVEC/fibroblasts), ADSC-EVs were visualised as being taken up by HUVECs, MDMs, and fibroblasts, demonstrated in Figure 4B-C. There was no obvious preferential uptake by any one cell type. This EV uptake was also evident when the models were visualised by 3D reconstruction using IMARIS software, where the red vesicles are observed colocalising with all cell types within the two developed models (Supplementary Data 1A-D).

**Figure 4:**
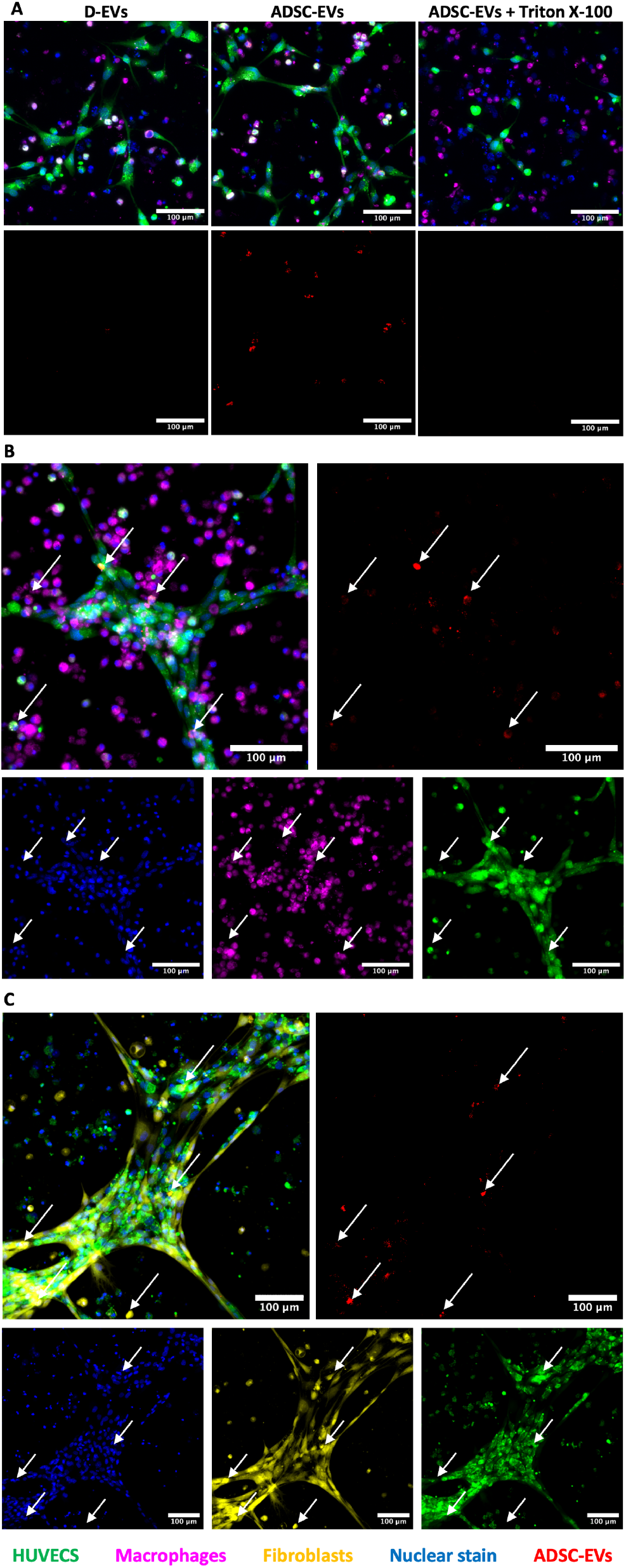
Optimisation of ADSC-EV staining and visualisation of EV-uptake by all cell types. A) D-EVs (left), ADSC-EVs (middle), or Triton X treated ADSC-EVs (right) were stained with PKH26 to confirm specific staining of ADSC-EVs with this dye (presence of red staining in middle panel). The top images are with all the channels turned on and the bottom are with only the ADSC-EV (red) channel on. EV uptake was observed in HUVEC/MDM models (B) and HUVEC/fibroblast models (C). White arrows indicate EV examples. Green = HUVECs, magenta = MDMs, yellow = fibroblasts, red = EVs, and blue = nuclear stain. Scale bar represents 100 μm.

### The developed models can be used to investigate functional changes in response to treatment with ADSC-EVs

3D multicellular models were cultured with the addition of ADSC-EVs or D-EVs to determine the functional impact of ADSC-EVs on HUVEC tube formation in the presence of other cell types. Compared to D-EVs (set to 1), cultures treated with ADSC-EVs had increased tube formation. With ADSC-EVs, all multicellular models demonstrated larger fold changes than with HUVECs cultured alone (Figure 5A-D). Relative to D-EVs added to HUVEC/MDM models, treatment with ADSC-EVs did not significantly increase any of the parameters measured, although there was a non-significant increase in both number of junctions and junction density. For HUVEC/fibroblast models, cultures treated with ADSC-EVs had significant average fold change increases in the number of junctions (2.60±0.51, *p*=0.0005, Figure 5A), vessel length (1.98±0.25, *p*<0.0001, Figure 5C), junction density (2.18±0.82, *p=*0.0264, Figure 5B), and vessel area percentage (1.98±0.79, *p=*0.0468, Figure 5D) compared to D-EVs. Representative images of the models are presented in Figure 5E (HUVEC single cell type model), Figure 5F (HUVECs/MDMs), Figure 5G (HUVECs/fibroblasts), and Figure 5H (full models). This increase in response to ADSC-EVs was further pronounced with the full models, which demonstrated significant increases in number of junctions (3.99±0.41, *p*<0.0001, Figure 5A), junction density (2.14±0.26, *p*=0.0322, Figure 5B), and vessel length (3.017±0.10, *p*<0.0001, Figure 5D).

**Figure 5:**
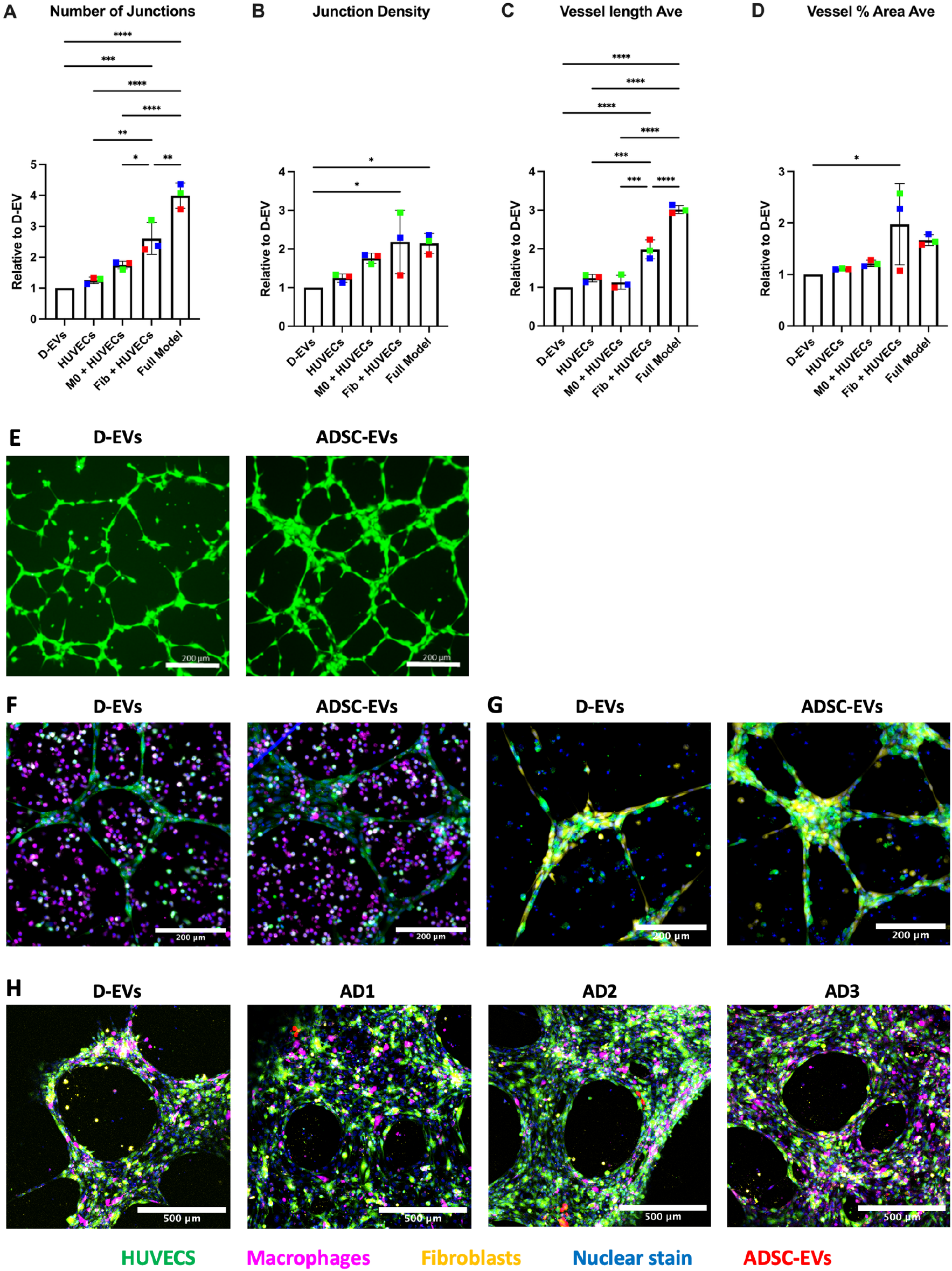
ADSC-EVs increase tube formation in 3D multicellular models. HUVECs were cultured alone, with MDMs, with fibroblasts, or all three cell types together, to assess tube formation. (A-D) ADSC-EVs increased tube formation across the different models when compared to D-EVs. Groups were compared using a one-way ANOVA and Tukey’s multiple comparisons tests. Coloured dots represent different ADSC patients (n=3) across 2 experimental runs. (E) Representative images of single cell type HUVEC cultures. (F, G Representative images of HUVEC/MDM and HUVEC/fibroblast co-cultures, respectively. (H) Representative images of full multicellular models treated with D-EVs, or EVs from three individual patients. Green = HUVECs, magenta = macrophages, yellow = fibroblasts, red = EVs, and blue = nuclear stain. Scale bar represents 200 μm (E-G) or 500 μm (H). Data were analysed using GraphPad Prism (9.1.0). \**p*<0.05, \*\**p*<0.01, \*\*\**p*<0.001, \*\*\*\**p*<0.0001.

3D multicellular models with the addition of ADSC-EVs or D-EVs were also studied to determine the functional impact on cellular co-localisation. A representative image of the analysis is demonstrated in Figure 6A-B. Compared to D-EVs, ADSC-EVs produced an increase in the proportion of MDMs localising with HUVEC tubes in the two-cell-type model (1.22±0.09, *p*=0.0152, Figure 6C). Fibroblasts and HUVECs were visualised to completely co-locate together within the models with D-EVs, and no change was observed following the addition of ADSC-EVs (Figure 6D).

**Figure 6:**
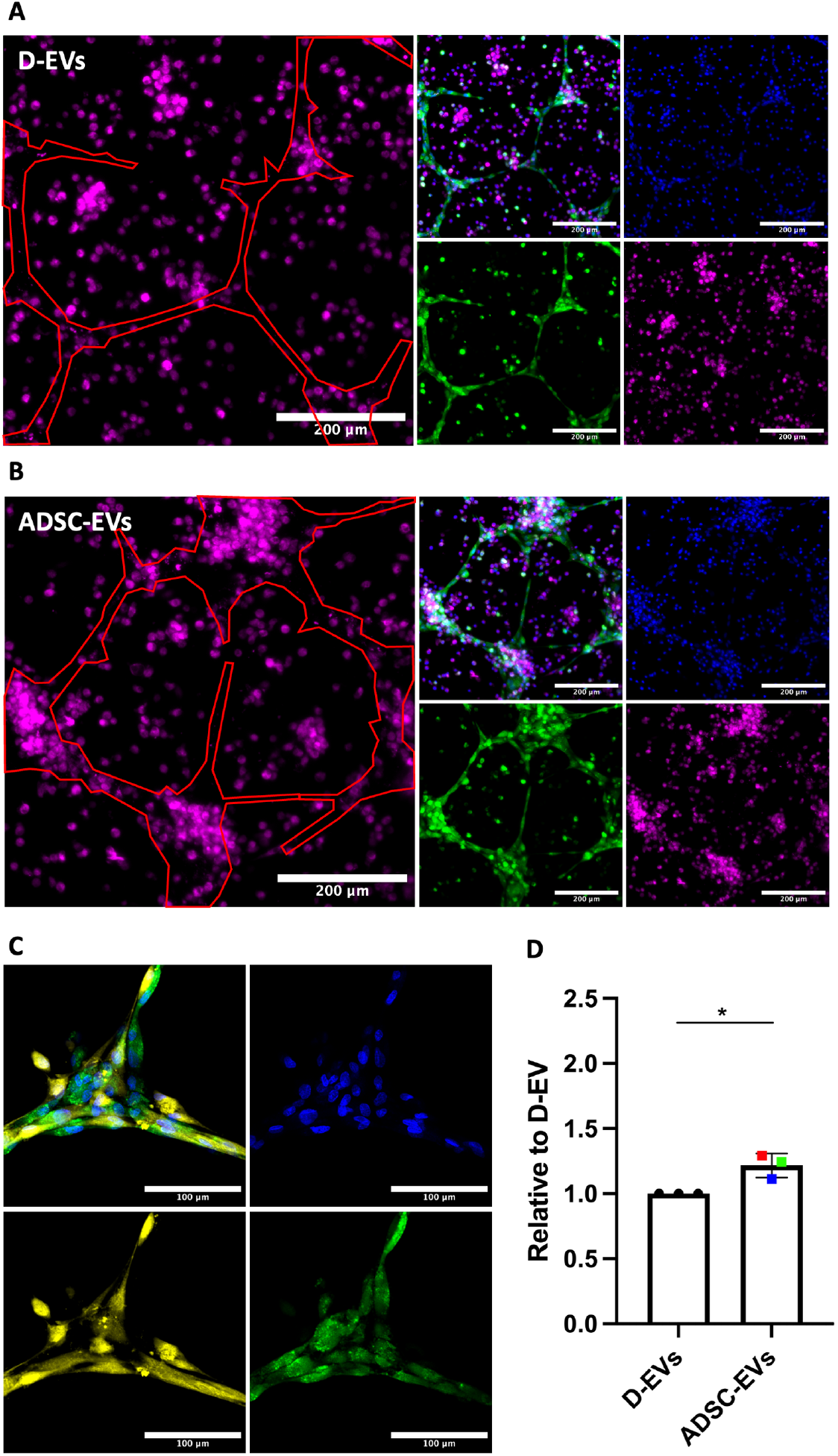
ADSC-EVs increase colocalisation of MDMs and HUVECs. HUVECs were co-cultured with either MDMs or fibroblasts to assess tube formation. Groups were compared using a t-test. Coloured dots represent different ADSC patients (n=3) across 2 experimental runs. \**p*<0.05. A-B) Representative images of co-localisation analysis traces. C) ADSC-EVs increase co-localisation of MDMs to HUVECs. Data were normalised to the HUVEC trace area (red) and compared using a one-way ANOVA. D) Representative image of fibroblasts and HUVECs with complete co-localisation. Green = HUVECs, magenta = MDMs, yellow = fibroblasts, red = EVs, and blue = nuclear stain. Scale bar represents 200 μm in A-B and 100 μm in D.

## Discussion

This study developed a 3D multi-cell-type *in vitro* model, which was then applied to study the effects of ADSC-EVs on vascularisation as a model relevant to wound healing. We observed that ADSC-EVs increased tube formation and MDM/HUVEC co-localisation, which was enhanced within the multicellular 3D models compared to single cell type models. This study is the first to investigate the functional role of ADSC-EVs on cells within multicellular *in vitro* systems.

Multiple cell types are involved in wound healing in an *in vivo* setting. We sought to better recapitulate this setting by focussing on the inclusion of fibroblasts and macrophages with endothelial cells to model vascularisation relevant to wound healing. Fibroblasts play an important role in wound healing as they provide the foundation for capillary networks to form around. This scaffold is vital for the successful development of new blood vessels to perfuse the injured area ^17^. The increase in tube formation that we observed when HUVECs were cultured with fibroblasts is expected as these cells coordinate to facilitate angiogenesis. When cultured together, fibroblasts and endothelial cells also completely co-localised and grew in structures around one another, consistent with the described biological functions of these cells to create capillary networks. A previous methods paper described a similar methodology to the current study for co-culturing fibroblasts and HUVECs to assess angiogenesis using Matrigel-coated plates ^18^. Several additional studies have investigated co-culturing these two cell types together in different matrices and models to assess angiogenesis. Sukmana and Vermette ^19^ previously co-cultured HUVECs and human dermal fibroblasts within a fibrin structure and concluded that fibroblasts enhance vessel formation as well as vessel development and maturation. This increase was even more pronounced when HUVEC cultures were supplemented with the growth factors vascular endothelial growth factor and basic fibroblast growth factor, both of which are known to stimulate angiogenesis. A further study by Liu *et al*. ^20^ co-cultured HUVECs and fibroblasts in a collagen structure, demonstrating that the presence of fibroblasts enhanced tube formation by 116%. While the matrices for cell growth differed across these studies, there has been consistency in findings for all of these fibroblast/HUVEC co-cultures, reinforcing the important role that fibroblasts play in facilitating angiogenesis. No previous study has investigated the role of ADSC- or MSC-EVs in these types of models. In adipose tissue, ADSCs are part of the stromal vascular fraction and form an important part of the heterogeneous collection of cells that makes up the adipose support system ^21^. Our study suggests that ADSC-EVs can further enhance the pro-angiogenic effects demonstrated in these co-culture models. As the first study to investigate ADSC-EVs in this context, we provide novel and important data to a largely understudied field that can progress research towards developing ADSC-EVs as potential therapeutics.

Macrophages also have an important role in wound healing and angiogenesis. When macrophages are activated during the wound healing process, they release growth factors, such as semaphorins and VEGF, that stimulate capillaries to start forming new sprouts, provide direction and guidance cues to establishing networks, and facilitate anastomosis ^22^. The presence of macrophages in this environment, therefore, directly impacts angiogenesis. We have previously concluded that ADSC-EVs reduce the overall inflammatory phenotype of macrophages ^8^, which may further promote angiogenesis in this context by providing a favourable (dampened inflammation) environment. It is therefore not surprising that, when cultured alongside macrophages, we observed an increase in HUVEC tube formation which was further enhanced with the addition of ADSC-EVs. In addition to increasing tube formation, ADSC-EVs increased the co-localisation of macrophages to HUVEC tubes, which is likely of further benefit to wound healing and tissue survival through direct cell-cell interactions. A study investigating the influence of mouse-derived macrophages on *in vitro* tube formation in a hydrogel structure found that M0 (such as those used in our study) and M2 (but not M1) macrophages enhance vessel formation and directly interact with endothelial cells ^23^. Furthermore, *in vivo* studies have demonstrated that macrophages co-locate with vessels to facilitate tube formation ^24^. To the best of our knowledge, we are the first to investigate these co-culture models using human cells. This enhances the relevance and translatability of our findings, especially in angiogenesis and wound healing, where species-specific differences can affect cellular responses ^25^. Therefore, while the mouse model data are not directly comparable to our human models, the findings reinforce the notion that macrophages directly influence angiogenesis.

Most studies involving 3D cell cultures focus on organoids, spheroids, or the use of bioengineered substances such as hydrogels. In this study, we aimed to develop 3D multicellular models that can be utilised and adapted by research groups using widely available materials and relatively simple techniques that could be adapted to higher throughput screens. The purpose of developing this model was for use as an *in vitro* tool for testing drugs or biologicals for their ability to aid vascularisation, tissue regeneration, or wound healing. In future, these models could be used for screening the ability of compounds to heal a scratch wound or to ‘rescue’ a tissue microenvironment that is in a state that is unfavourable for wound repair (e.g. through the addition of inflammatory cytokines or other biological agents).

Matrigel is a basement membrane substance routinely used in cell culture experiments with well-established protocols for its use. Our first optimisation step was to determine the most functionally relevant way to seed each cell type into this model. Method 1, where both cell types were combined in Matrigel prior to plating them, did not result in successful tube formation. This method was initially chosen as *in vitro* studies using hydrogels are developed by first mixing the cell suspensions with the gelation substance before polymerisation ^26^. While this method works within a hydrogel structure, it did not transfer to our preferred Matrigel system. HUVECs form tubes best when they are seeded on top of pre-set Matrigel as they can naturally enter the structure and form tubes. For this reason, we next attempted mixing the macrophages in Matrigel prior to seeding HUVECs (in media) on top (Method 2). While tubes formed, the structures were very thick (∼300 μm), and the macrophages were too sparse to interact with the HUVECs. As the HUVECs were seeded on top, they did not penetrate throughout the entire 3D structure, reducing their potential to interact with macrophages that were seeded evenly throughout the structure. This is unlikely to be reflective of processes *in vivo*, as paracrine signalling is fundamental in both the initiation and control of inflammation ^27^. If cells are too sparse, paracrine signalling is hindered, and normal wound healing would be impeded due to lack of cell proximity. We determined that seeding both cell types on top of Matrigel (Method 3) allowed for both tube formation and co-localisation of these cells.

Further investigations also demonstrated that fibroblasts and HUVECs interact very strongly when seeded into a 3D multicellular model in this way. This model is also representative of how macrophages enter the tissue. In response to tissue injury or damage, monocytes travel through the blood to the site of the injury, where they differentiate into macrophages and facilitate inflammation ^28^. By creating a 3D layer that encompasses multiple cell types interacting with one another, we can mimic the wound microenvironment in a way that can be manipulated to investigate wound healing in different tissue microenvironments in the future.

We demonstrated that ADSC-EVs significantly enhanced HUVEC tube formation, and this was observed to a greater extent when cultured in multi-cell-type models with macrophages and fibroblasts, compared to when HUVECs were 3D cultured alone. Furthermore, when examining the control group (D-EVs), we also observed an increase in tube formation within the multicellular models compared to the HUVECs cultured alone. This highlights the importance of using more complex multicellular models rather than relying solely on single-cell cultures when evaluating potential therapeutic options, as these more complex models better mimic the interactions and dynamics of the *in vivo* microenvironment. Culturing different cell types together accounts for cell-cell communication and interactions that influence cellular functions. Angiogenesis is a vital process for wound healing as capillary development provides nutrients and oxygen to the damaged tissue to promote tissue survival and prevent necrosis ^1,29^. Endothelial cells are stimulated and work in conjunction with other cells during the capillary sprouting process. It was therefore important to investigate the effect on angiogenesis in these newly developed multicellular models.

Our multicellular models for assessing angiogenesis in wound healing can be adapted to study various tissue structures and disease pathologies. Future work could substitute different cell types within the models to replicate specific microenvironments depending on the tissue structure or pathology of interest. For example, the addition of cancer cells to this model could be used to investigate cell invasion and metastasis. There are, however, limitations to the model design; these include the number of cell types that can physically be incorporated into the models, as well as the type of microscope as a limiting factor for how many fluorescent stains can be incorporated into each panel. These factors would need to be taken into consideration when developing future models of complex tissue microenvironments.

In this study, we have developed a multicellular model that can be used for future work assessing wound healing *in vitro*. Compared to single cell type models, our model more effectively captures the complexity of the wound microenvironment by incorporating multiple cell types in a 3D structure. This more accurately reflects the dynamic nature of wound healing, which relies on intricate interactions between various cell populations. Furthermore, using our models, we have shown that ADSC-EVs significantly enhance tube formation in HUVECs. These findings add to a body of evidence that highlights the potential of ADSC-EVs as a promising therapeutic approach for improving wound healing and tissue regeneration.

## Acknowledgements

We offer special thanks to the Hugh Green Foundation for their ongoing support and funding of the Hugh Green Technology Centre – Bioimaging Core at the Malaghan Institute of Medical Research provided expertise on image acquisition and analysis.

